# A Bioelectrochemical Crossbar Architecture Screening Platform (BiCASP) for Extracellular Electron Transfer

**DOI:** 10.1101/2025.07.09.663982

**Authors:** Hasika Suresh, Presley Bird, Kundan Saha, Surya Varchasvi Devaraj, Cihan Asci, Albert Truong, Cooper Wallman, Rhea Patel, Atul Sharma, Matthew D. Carpenter, Sujitkumar Bontapalle, Rafael Verduzco, Jonathan J. Silberg, Sameer Sonkusale

## Abstract

Electroactive microbes can be used as components in electrical devices to leverage their unique behavior for biotechnology, but they remain challenging to engineer because the bioelectrochemical systems (BES) used for characterization are low-throughput. To overcome this challenge, we describe the development of the Bioelectrochemical Crossbar Architecture Screening Platform (BiCASP), which allows for samples to be arrayed and characterized in individually addressable microwells. This device reliably reports on the current generated by electroactive bacteria on the minute time scale, decreasing the time for data acquisition by several orders of magnitude compared to conventional BES. Also, this device increased the throughput of screening engineered biological components in cells, quickly identifying mutants of the membrane protein wire MtrA in *Shewanella oneidensis* that retain the ability to support extracellular electron transfer (EET). BiCASP is expected to enable the design of new components for bioelectronics by supporting directed evolution of electroactive proteins.

**The bigger picture:** Devices that interface microbes and materials, known as bioelectronics, can be used to sense environmental chemicals in real time, generate energy from sugars, and synthesize chemicals. While these devices leverage the unique capabilities of living systems as components in devices, such as their ability to convert chemical information in the environment into electrical information at the cell surface, it remains challenging to engineer these cellular components and their biomolecules for new applications, largely because commercially available bioelectrochemical systems for monitoring current generated by electroactive microbes are costly and require large culture volumes, needs continuous monitoring for days to obtain stable signals, and multichannel potentiostats to monitor multiple microbes in parallel.

To overcome these challenges, we created the Bioelectrochemical Crossbar Architecture Screening Platform or BiCASP that is easy to fabricate, enables parallel analysis of microbial samples in flexible arrayed formats, and yields a stable signal on the minute time scale. This device is expected to enable the application of combinatorial protein engineering methods, such as directed evolution, to proteins that control microbial current production, by allowing for fast screening of cells expressing protein mutant libraries. As a proof-of-concept, we demonstrate that this device can screen for cells that express mutants of decaheme cytochromes that retain the ability to electrically connect cells to electrodes. This device will simplify the engineering of cells and proteins that function as electrical switches as well as the diversification of bioelectronic devices for real-time sensing of chemicals in the environment.

Furthermore, BiCASP is promising as a high-throughput screening (HTS) platform, enabling rapid, parallel analysis of cellular and molecular interactions of diverse biological systems through label-free electrochemical methods. Such capabilities could transform drug discovery, personalized medicine, and functional genomics, supporting systematic genetic and chemical screens even at single-cell resolution.

**Highlights:**

- A high-throughput screening platform with individual addressability
- A device with a flexible crossbar architecture that simplifies current analysis
- Reproducible detection of real-time cellular current on the minute time scale
- The device can be used to screen a library for cells with functional protein wires

**Graphical Abstract:** 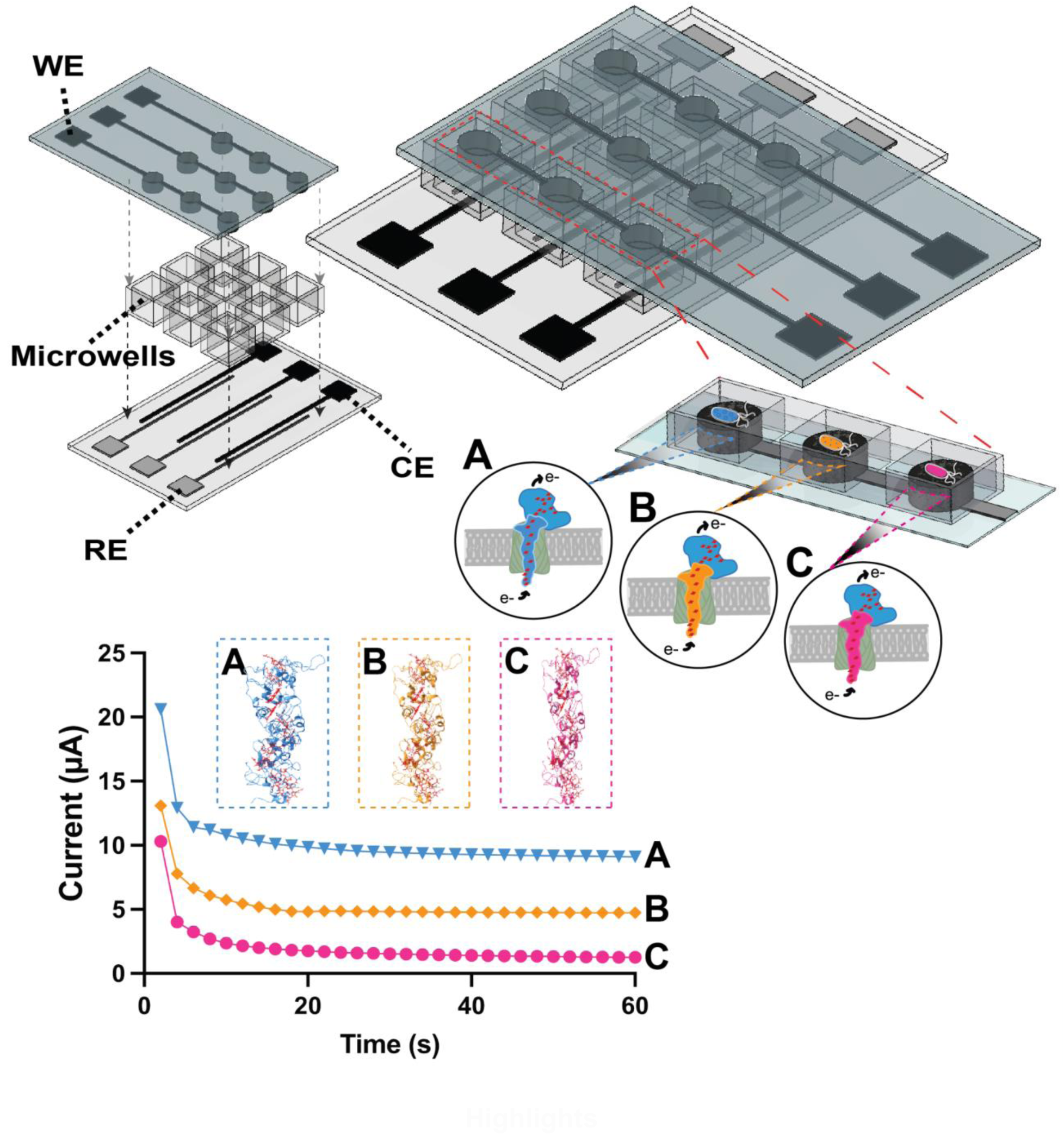

## Introduction

High-throughput screening (HTS) platforms are essential in modern biology and life sciences, enabling rapid, parallel investigation of cellular responses, signaling pathways, and biomolecular interactions.^1^ These platforms have transformed drug discovery,^2^ personalized medicine,^3^ and functional genomics^4^ by allowing systematic screening of genetic perturbations and chemical libraries, with some platforms achieving single-cell resolution.^5^ Among emerging HTS technologies, electrochemical approaches offer distinct advantages: label-free, real-time, and highly miniaturizable detection with low sample volume and low power requirements. ^6–8^ These platforms have been widely used for systems biology to monitor cell metabolism,^9^ redox states,^10^ ion transport,^11^ and secreted analytes,^12^ and are increasingly integrated with microfluidic and organ-on-chip systems for multiplexed, high-resolution assays.^13^ While electrochemical HTS platforms have been diversified for systems biology, they have not been leveraged to the same extent for synthetic biology, where scalable, real-time monitoring of engineered cellular function is needed for pathway optimization, protein engineering, and strain development.^14^

Microbes can be used as electrical components of devices to sense chemicals,^15–17^ produce energy from organic matter,^18,19^ support remediation of pollutants and waste,^20,21^ and perform electrosynthesis.^22,23^ In these bioelectrochemical systems (BES), microorganisms on the surface of an electrode couple biochemical reactions within cells to electrode oxidation and reduction.^24–28^ These microbes accomplish this coupling through a process called extracellular electron transfer (EET), which is often facilitated by transmembrane wires^29,30^ such as the Mtr protein complex.^31^ Electroactive microbes having the Mtr complex are found in diverse environmental niches.^32,33^ However, it remains arduous to continuously monitor the current generated by these electroactive microbes and to obtain stable and reproducible signals. The BESs used to perform chronoamperometry are low throughput, requiring reactors having large culture volumes (≥10 mL), constant growth medium input, and continuous monitoring for long durations (>1000 minutes) to achieve stable signals. These systems are also expensive, limiting accessibility for many researchers.

One way to enable new applications of bioelectronic devices is to engineer conductive proteins with new sequences and functions^34,35^ and then to express those proteins within electroactive microbes interfaced with electrodes.^16^ By targeting oxidoreductases for protein design, cytosolic protein electron carriers have been designed that exhibit conditional electron transfer^36–39^ and altered electrochemical properties.^40,41^ However, the low throughput of BES measurements with electroactive microbes has severely limited our ability to engineer transmembrane protein wires that mediate EET, such as MtrA, using combinatorial approaches like directed evolution,^42,43^ where libraries of biomolecules and cells are screened for specific functions using high-throughput assays.^44–47^

To date, only a handful of mutations in the Mtr pathway have been evaluated using BESs.^48,49^ While strategies have been identified to increase the throughput of EET measurements by performing high throughput growth selections,^50^ these approaches only provide a snapshot on the total current generated by cells over long incubations rather than real-time information about the current that electroactive microbes produce on an electrode surface. To increase the throughput of dynamic current measurements with electroactive microbes under the conditions where bioelectronic technologies are being performed, we developed a <u>Bi</u>oelectrochemical <u>C</u>rossbar <u>A</u>rchitecture <u>S</u>creening <u>P</u>latform (BiCASP), which can be fabricated with a flexible number of sample wells at a cost of ≤$15 per device. Prior work on HTS of electrogenic bacteria is limited, largely implemented on printed circuit boards (PCBs) and paper, primarily to screen for bacteria suitable for energy harvesting from wastewater using microbial fuel cells (MFC).^51,52^ Using a 24-well BiCASP, we show how this device allows for facile biological workflows beyond MFCs, where the real-time current generated by electroactive microbes can be rapidly characterized in a three-electrode configuration. By comparing *Shewanella oneidensis* MR-1 containing and lacking the Mtr pathway,^53^ we show that BiCASP can differentiate cells that present EET from those that do not on the minute time scale in sample wells containing small volumes (<1 mL). As a proof-of-concept application to illustrate the utility of this system, we use the BiCASP to screen a library of MtrA peptide insertion mutants for variants that support EET in *S. oneidensis* MR-1, and find that this device can be used to rapidly discover mutants that support EET, like MtrA.

## Results and Discussion

### The Bioelectrochemical Crossbar Architecture Screening Platform (BiCASP)

Bioelectrochemical Systems (BES) are regarded as the gold standard for measuring direct EET in exoelectrogens.^54^ However, these systems are limited to measuring EET of only a single strain at a time, resulting in low throughput and increased processing times. This makes these systems incompatible with HTS of protein mutant libraries, which is needed to leverage directed evolution for protein design.^42,43^ To screen libraries of decaheme protein mutants produced by protein engineering for EET, we propose a high-throughput system with individually addressable bioelectrochemical microwells that monitor EET activity in parallel (Figure 1). *<u>This device is designed to increase the throughput of electroactive microbe characterization by allowing for parallel measurement of current from multiple strains, or strains expressing different mutant proteins, within minutes, as opposed to hours or days</u>*.

**Figure 1.**
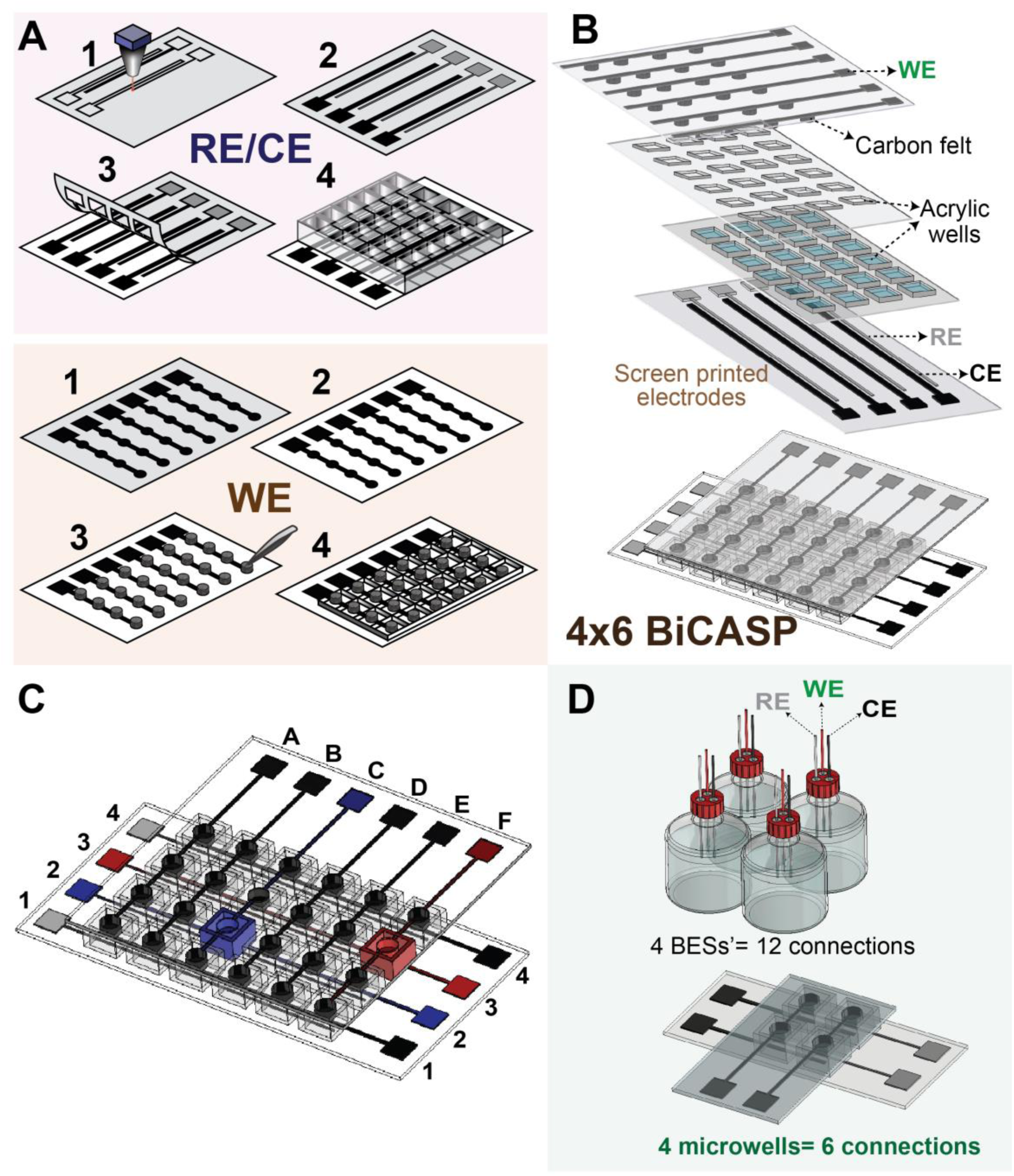
An overview showing the BiCASP architecture and connections. (**A**) Schematic showing the fabrication steps to make the BiCASP. For the CE/RE, Step 1: The stencil design is cut with a laser cutter, Step 2: The stencil is pasted onto a polyethylene terephthalate (PET) sheet and the electrodes are screen printed, Step 3: The stencil is peeled off exposing the RE and CE, Step 4: Acrylic well (1/4^th^ inch) are attached to the PET sheet. For the WE, Step 1: The stencil design is cut with a laser cutter, Step 2: The stencil is pasted onto a PET sheet and the WE is screen printed, Step 3: Carbon felt is attached to the circles on the WE, Step 4: Acrylic well (1/16^th^ inch) are attached to the PET sheet. **(B)** A deconstructed and an assembled version of the BiCASP in a 4x6 configuration. The RE and CE are on the bottom plane with acrylic wells housed right above each electrode pair, while the WE is used in the crossbar architecture from the top plane. (**C**) Image showing how individual addressability of cells can be achieved with a 4x6 BiCASP. (**D**) Comparison images of BiCASP in a 2x2 configuration showing the reduction in the number of connections needed for the same number of wells/cells and offering edge readout for easy handling and operation.

BiCASP incorporates elements of a traditional three-electrode system arranged in two distinct planes, resulting in a three-dimensional configuration. These are essentially miniaturized single-chamber BES. The fabrication steps to make the BiCASP are shown in Figure 1A. The reference electrode (RE) and the counter electrode (CE) lie on the bottom plane, while the working electrode (WE) is in a crossbar architecture on the top plane (Figure 1B). Carbon felt is used as the WE because of its porous, conductive, and biocompatible nature.^55^ Silver-silver chloride ink is used as the RE, while conductive carbon ink is used as the CE. Patterned acrylic sheets function as microwells to contain media and bacteria. Each microwell measures 1 cm^2^ and has a capacity of 500 µL. The BiCASP can be customized for any number of wells (9, 16, 24, 96, and 384 wells) to be compatible with different high-throughput platforms used for cellular growth (Figure S1). For this study, we used a 24-well system to conduct all experiments. This architecture was chosen to align with biological workflows used for growing arrayed cellular samples. This planar crossbar architecture of BiCASP offers significant advantages in scalability and density, enabling the vertical stacking of electrodes and facilitating 3D integration, which is challenging with wired electrodes on a single plane.^56^ *<u>This architecture offers decreased signal propagation distances and parallel operation capabilities within different arrayed formats that can be aligned with the formats used in biology to prepare samples of electroactive microbes</u>*.

Each well is individually addressable due to the BiCASP configuration. In a given column (Figure 1C), all wells share a RE and a CE. The rows share the same WE. Due to this orthogonal positioning of the RE/CE and WE, a single microwell can easily be selected for analysis without the need for electronic switches. To activate a microwell at the location (X, Y) in this two-dimensional array, the RE line on column Y, the WE line from row X, and the CE line from column Y are chosen to be connected to the potentiostat. For instance, as illustrated in Figure 1C, connecting RE2/CE2 and WE(C) to the potentiostat solely activates the microwell at location (2,C) shown in blue, whereas connecting RE3/CE3 and WE(F) activates only the dark red well at location (3,F). *<u>With this design, the individual addressability no longer requires dedicated selection logic and electronic circuitry. The complexity of the interconnect (wiring) is easily resolved due to the crossbar architecture of BICASP, and the platform can be scaled to large arrays</u>*.

In a prior study, a 96-well electrochemical microwell plate (ec-MP) was developed for high-throughput screening of EET generated by electroactive microorganisms.^57^ However, each well was equipped with three electrodes, which resulted in a total of 96 x 3 = 288 complex interconnect wiring connections to be connected to the readout electrochemical unit.^57^ In contrast, a BiCASP with the same number of wells has a decreased number of connections because it utilizes shared reference and counter electrodes on one plane, and an orthogonal working electrode (WE) on a different plane (Figure 1D). *<u>In this way, BiCASP requires only 8 reference, 8 counter and 12 working electrodes for a total of just 28 interconnect/wirings to access 96 individual microwells</u>*.

With BiCASP, the connection pads located at the end of the electrode line simplify the process of connecting the well(s) and sample(s) selected for analysis to the potentiostat. To enhance the automation of the BiCASP, we incorporated an optional electrical multiplexer (MUX) to programmatically scan through the array. This system comprises three 8x1 multiplexers – one for each electrode – along with select lines, signal lines, and a grounding port (Figure S2A). Each MUX is responsible for selecting one CE, WE, and RE at a time, thereby connecting the respective electrodes to a specific well within the BiCASP array. By using 12 control lines, the microcontroller (Figure S2B) facilitates the switching between any combination of RE, WE, and CE at a desired switching frequency, thereby enabling the automatic operation of the BiCASP. We developed a user interface to configure the MUX parameters, such as switching frequency, well selection, and the number of cycles (Figure S2C), to achieve full automation. *<u>Overall, the BiCASP is a miniaturized three-electrode BES capable of measuring EET in multiple strains, or single strains with different mutations, in a highly parallelized and automated manner</u>*.

### Identifying optimal conditions for BiCASP detection of EET

To first evaluate the electrochemical performance of the carbon felt in the BiCASP, we performed cyclic voltammetry (CV). When KCl (0.1 M) was added to a BiCASP well (Figure 2A), there was no distinct redox signal. In contrast, the addition of a known redox couple – Ferro/Ferricyanide (1 mM) resulted in oxidation (0.27 V) and reduction (0.05 V) peaks similar to previously reported values.^58^ *<u>Together, these results indicate that carbon felt can serve as an effective working electrode</u>*.

**Figure 2.**
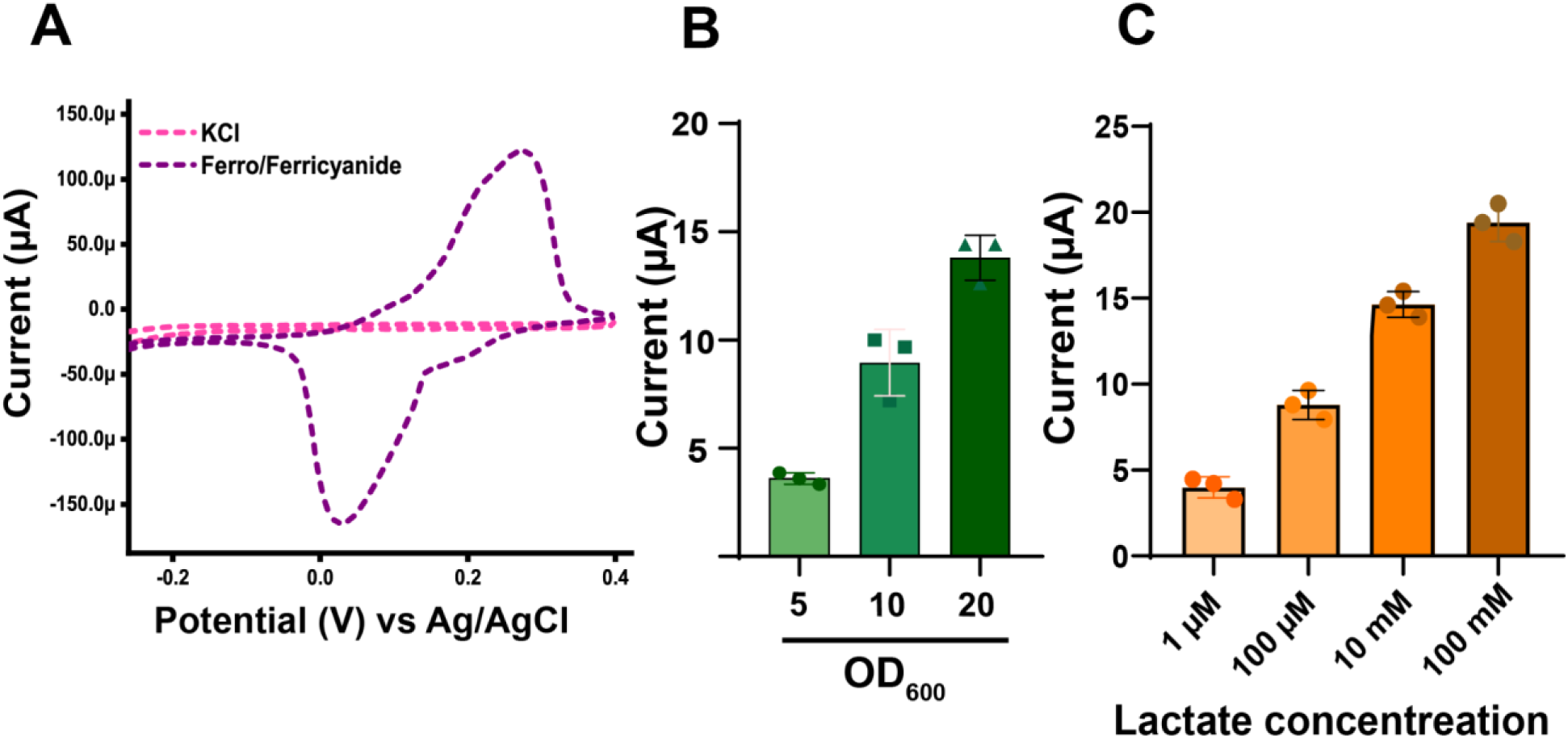
Identifying BiCASP conditions that yield strong biological signals. (**A**) CV curves of 0.1 M KCl and 1 mM ferro/ferricyanide showing the device’s sensitivity for redox active species. Chronoamperometry at 0.2 V vs. Ag/AgCl with MR-1 at (**B**) different cell concentrations (OD_600_ refers to the optical density (600 nm) of a 50 µL suspension of cells added to the microwell) (Linear regression analysis; R^2^ = 0.91 p <0.0001) (**C**) different lactate concentrations (Linear regression analysis; R^2^ = 0.66, p = 0.0014). The current increases with increasing bacterial load and lactate concentration in both cases. Bars represent the average current observed 180 seconds after beginning measurements, calculated from three biological replicates. Error bars show ±1σ.

To establish optimal operational parameters for using BiCASP to detect EET, we initially measured the current arising from adding various concentrations of *S. oneidensis* MR-1 to the WE. When the current was measured in M9 minimal medium containing 50 mM sodium lactate, a stable current was observed for each concentration within 180 seconds (Figure S3A), compared to 15 - 30 hours typically needed in a large microbial BES.^24–28^ The magnitude of the current generated was proportional to the number of bacterial cells added to the WE (Figure 2B). To understand how current is affected by the amount of lactate provided, which represents the carbon source that *S. oneidensis* MR-1 uses for growth and EET,^59^ current was measured across a range of lactate concentrations using 50 µl of bacterial culture, OD_600_ = 20. With these measurements, the current was monotonic and positive to the lactate concentration (Figures 2C, S3B). *<u>These findings demonstrate that the current observed in the BiCASP is dependent upon both the amount of bacteria added to the WE and the concentration of lactate added to the well. To ensure a reliable current when using BiCASP to study EET in samples, all subsequent measurements applied cells (50 µL) at an OD_600_ of 20 to the WE and used liquid growth medium having 100 mM lactate</u>*.

### BiCASP detects decaheme complementation of EET in bacteria

Existing approaches for monitoring microbial EET acquire data on the hour-to-day time scales, such as cell growth,^50^ BES measurements,^16^ iron reduction, and nanoparticle reduction.^60^ To investigate whether the BiCASP can be used to report on direct EET by acquiring data on the minute time scale, we compared the current generated by the electroactive microbe *Shewanella oneidensis* MR-1 and a mutant strain (JG665) that lacks genes required for EET (Figure 3A). With all BiCASP measurements, the current stabilized within 30 seconds of initiating measurements (Figure S4). After one minute, the average current presented by *S. oneidensis* MR-1 was significantly higher (7.5-fold) than that of *S. oneidensis* JG665 (Figure 3B). The coefficient of variance for the current produced by *S. oneidensis* MR-1 was 16%, while the coefficient of variance for *S. oneidensis* JG665 was 44%. Analysis of EET using an Fe(III) reduction assay revealed similar trends (Figure 3C), although the time required for signal detection was >100-fold longer (Figure S5). *<u>These findings show that BiCASP can reliably detect microbial EET mediated by multiheme cytochromes using short, minute-scale measurements</u>*.

**Figure 3.**
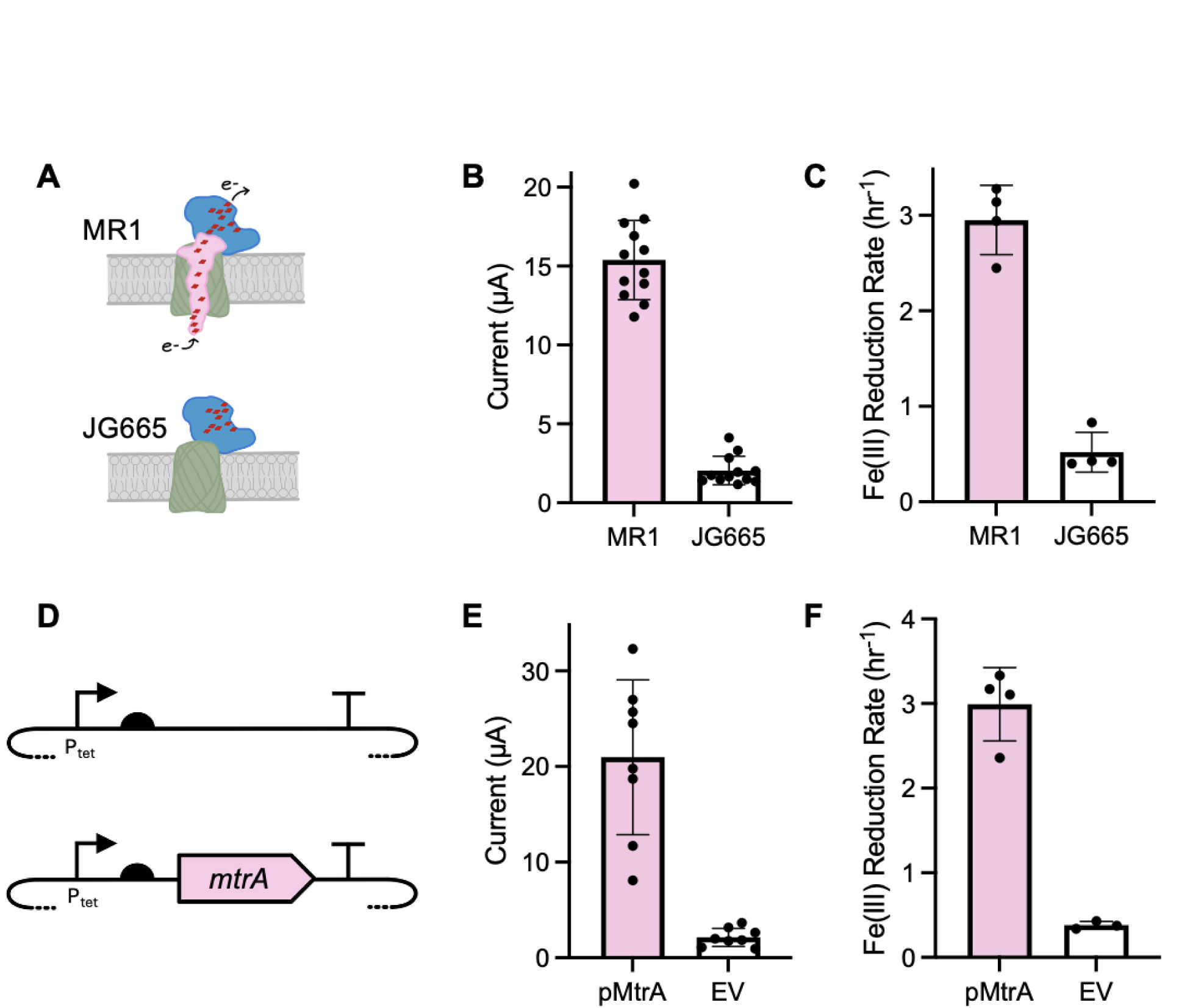
BiCASP can differentiate bacteria that exhibit EET from those with EET-inactivating mutations. (**A**) Two strains were compared using BiCASP, including *S. oneidensis* MR-1 and *S. oneidensis* JG665; the latter lacks *mtrA* and cannot support EET.^53^ (**B**) *S. oneidensis* MR-1 yields current on BiCASP that is significantly higher than that observed with *S. oneidensis* JG665 (Welch’s t test, p < 0.0001). (**C**) Visual detection of Fe(III) reduction using a ferrozine assay reveals similar differences (Welch’s t test, p = 0.0001). (**D**) A plasmid that constitutively expresses MtrA and an empty vector were used to evaluate complementation of *S. oneidensis* JG665 EET. (**E**) *S. oneidensis* JG665 transformed with the MtrA plasmid present a significantly higher current than cells harboring the empty vector (EV) (Welch’s t test, p = 0.0003). (**F**) Iron reduction measurements reveal similar trends (Welch’s t test, p = 0.001). For BiCASP measurements, the data represent the signal sixty seconds after beginning measurements. Data are presented as the average of eight or more biological replicates with error bars representing ±1σ.

With *S*. *oneidensis* JG665, EET can be restored by transforming cells with a plasmid that expresses the decaheme cytochrome MtrA.^53^ To investigate if BiCASP can be used to screen for plasmid complementation of *S. oneidensis* JG665 EET, we compared the current generated by *S. oneidensis* JG665 harboring a plasmid that constitutively expresses MtrA and cells containing an empty vector (Figure 3D). Cells transformed with the MtrA vector presented current after one minute, which was significantly higher (12.1-fold) than that of cells containing the empty vector (Figure 3E). The signal with cells expressing MtrA was slightly higher (1.4-fold) than that observed with *S. oneidensis* MR-1, although greater variability (coefficient of variance of 38%) was observed with MtrA complementation of EET. Analysis of each sample using the iron reduction assay revealed similar trends as the BiCASP (Figure 3F), albeit requiring a longer time scale for signal detection. *<u>These findings show that BiCASP can reliably detect plasmid complementation of an EET-deficient strain using short, minute-scale measurements</u>*.

### BiCASP can screen an MtrA mutant library for variants that mediate EET

In a prior study, a library of vectors was created that expresses MtrA variants having the octapeptide SGRPGSLS randomly inserted at different locations.^50^ The activity of these MtrA mutants was evaluated by monitoring the growth complementation of *S. oneidensis* JG665,^50^ an indirect measure of EET. While this study comprehensively mapped the ability of >200 mutants to support growth dependent upon EET, current was only measured for cells expressing 4 different MtrA mutants, due to the low throughput of BES measurements. To first investigate if BiCASP can be used to screen this library of MtrA mutants for variants that support EET (Figure 4A), we transformed *S. oneidensis* JG665 with this plasmid library and enriched for cells that complement growth under conditions that require EET. Following selection, cells harboring MtrA plasmids were spread on agar plates, and individual colonies were used to grow cultures for BiCASP analysis.

**Figure 4.**
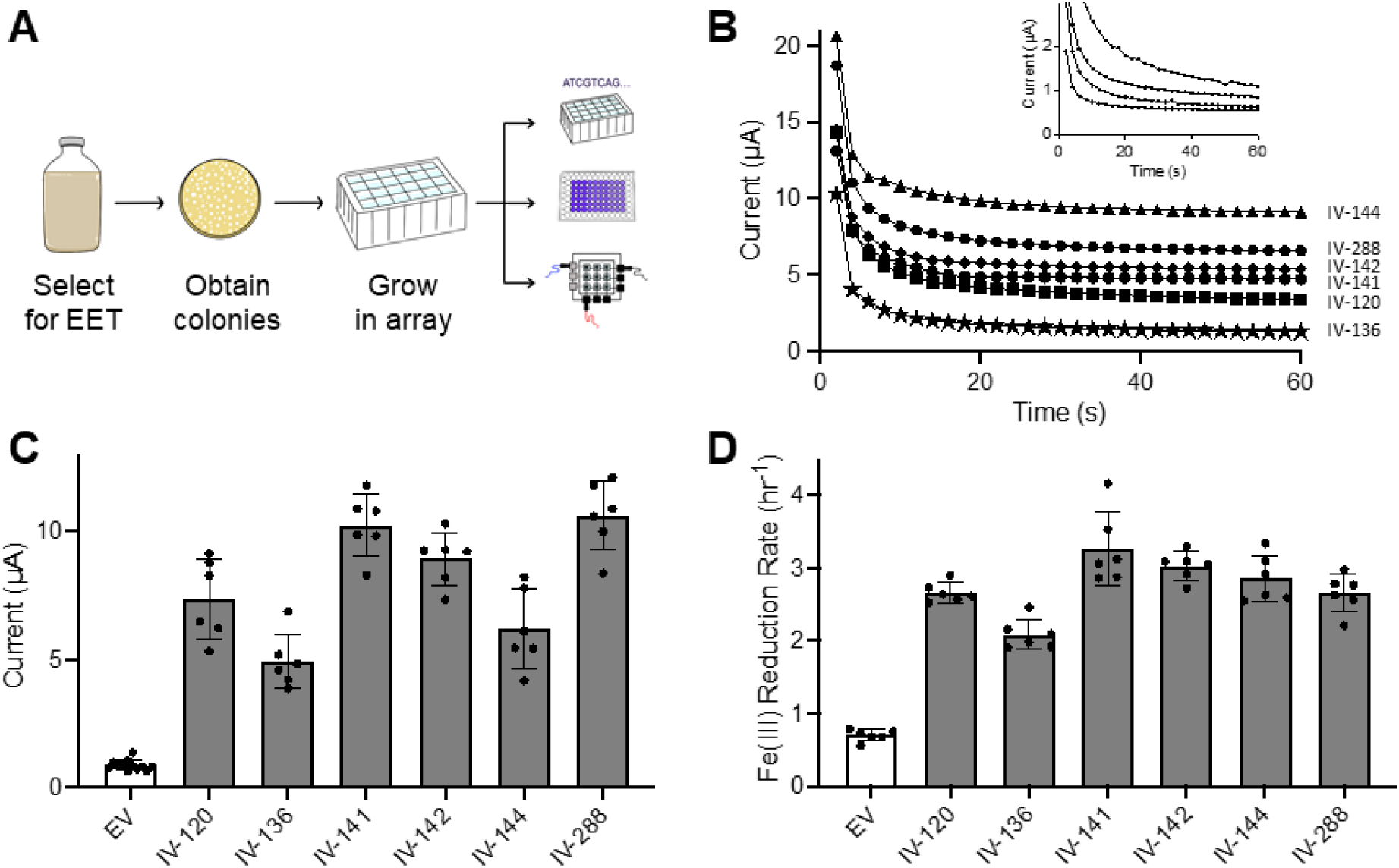
BiCASP reliably identifies active MtrA mutants selected for EET. (**A**) The library of MtrA mutants was grown anaerobically in medium containing ferric citrate as the only terminal electron acceptor to select for cells expressing MtrA mutants that support EET. Individual colonies were obtained by plating this culture on LB-agar medium, and these colonies were characterized using BiCASP, the Fe(III) reduction assay, and plasmid sequencing. (**B**) Current generated by a single replicate of cells derived from the different colonies were compared with *S. oneidensis* JG665 transformed with an empty vector. Each insertion variant (IV) is named based on the location of the peptide insertion. JG665 data is shown in the graph inlay. (**C**) BiCASP analysis of six biological replicates for each sample. Current represents the signal sixty seconds after beginning the measurement. With all insertion variants, the signal was significantly higher than in cells containing an empty vector (EV) (Welch’s t test, p ≤ 0.0004). Twelve biological replicates were analyzed for the negative control. (D) Iron reduction rates for the same samples revealed each insertion variant presents a signal that is significantly higher than the EV (Welch’s t test, p < 0.0001). Bars represent the average, while error bars show ±1σ.

When six different isolates were screened for EET using BiCASP (Figure 4B), five presented signals from a single measurement that were >10 σ out from the negative control (10.8 to 34.9 σ); one variant was only 2σ higher. To investigate if the EET signal observed when screening is reproducible, multiple biological replicates of each variant were performed using BiCASP. The average current for all six variants was significantly higher than that of the negative control (Figure 4C). Similarly, all six variants presented MtrA-dependent EET with the iron reduction assay (Figure 4D). *<u>These results show that single measurements of cells expressing different MtrA mutants can be reliably differentiated from the negative controls, illustrating how BiCASP can be used to mine protein mutant libraries for mutants that support EET</u>*.

To investigate if BiCASP can screen the peptide insertion library for MtrA variants that support EET without prior enrichment, *S. oneidensis* JG665 harboring the plasmid library was grown on LB-agar medium in the absence of selective pressure for EET. Individual colonies grown overnight were analyzed for EET using the BiCASP. In addition, cells were analyzed for iron reduction, and the plasmids in each culture were purified and sequenced to determine which MtrA mutants were expressed in each sample. BiCASP screening using two different 24-well devices revealed that MtrA insertion variants presented currents that varied from 2.6 to 12.9 µA (Figure 5A). Two variants had currents that were within 2σ of the negative control, while the remaining variants (94%) presented currents that were 2.3 to 12.1 σ higher than the negative control. Additionally, three variants (IV-282, IV-287, and IV-297) were observed in two different arrayed samples; the wells containing identical MtrA mutants presented similar currents. Iron reduction measurements revealed that 86% of the insertion variants exhibited activity that were >2σ higher than cells lacking MtrA (Figure 5B). These results show that a majority of the cells that present EET using BiCASP also present a signal in the iron reduction assay, although more subtle EET differences were observed between samples with BiCASP compared to the iron reduction assay.

**Figure 5.**
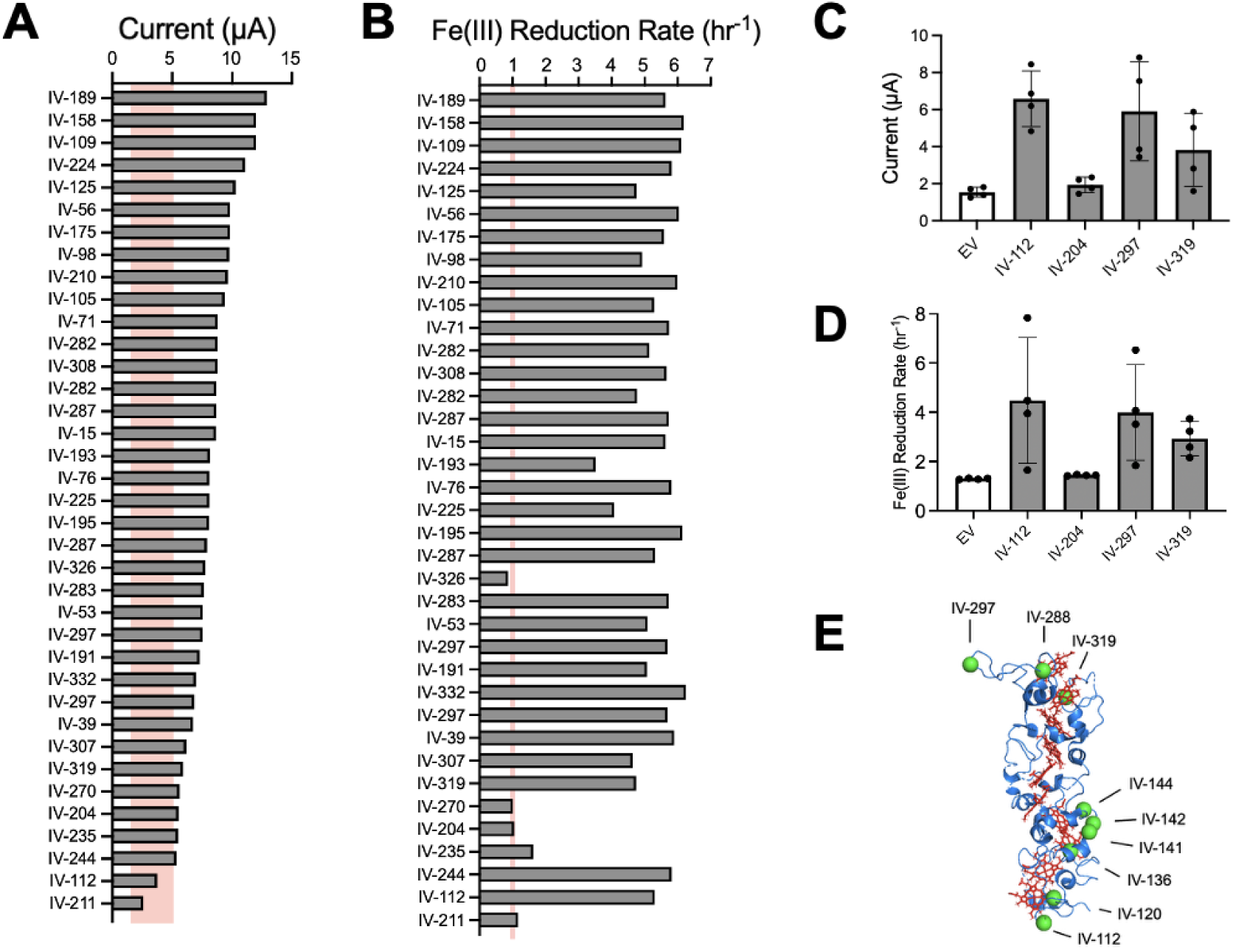
BiCASP increases the throughput of MtrA mutant library screening. Individual strains from a library of *S. oneidensis* JG665 harboring vectors that express different MtrA mutants were screened using (**A**) BiCASP and (**B**) the iron reduction assay. In total, 40 unique colonies were analyzed, but only those containing a peptide insertion are shown (n = 37). All data represent a single biological replicate, while the red line represents the average signal ±2σ obtained from cells harboring an empty vector. Four insertion mutants presenting low current from the initial screen were evaluated using larger numbers of biological replicates (n = 4) using (**C**) BiCASP and (**D**) the iron reduction assay. With BiCASP, IV-112 and IV-297 were significantly higher than the empty vector (EV) control (one tailed Welch’s t test, p < 0.05), while IV-319 had a p value of 0.051). In the iron reduction assay, the signals for all variants presenting more than 2x higher iron reduction rates were significantly higher than the empty vector (EV) control. (**E**) MtrA retains EET when peptides are inserted at locations spanning the outer membrane. The insertion sites of peptides (green spheres) are shown for variants whose EET activity was validated using multiple biological replicates on BiCASP.

To better understand the quality of the signal with variants that present low current using BiCASP, four samples were further characterized using a larger number of replicates. Specifically, we characterized IV-112 which lacked a signal on BiCASP but presented EET in the iron-reduction assay, IV-204 which lacked a signal with the iron reduction assay but presented a low current on BiCASP that exceeded the negative control, and and two variants (IV-297 and IV-319) that presented strong signals with the iron reduction assay and low signals on BiCASP. We found that IV-112, IV-297, and IV-319 all presented EET on BiCASP that was 2.5 to 4.3-fold higher than the negative control (Figure 5C), while IV-204 was only 1.25-fold higher. Among these variants, only IV-112 and IV-297 presented signals that were significantly higher than the negative control (p < 0.05); the p value for IV-319 was 0.051. Similar trends were observed when the iron-reduction assay was used to perform larger numbers of replicates (Figure 5D). However, with this assay, IV-112, IV-204, and IV-319 all presented rates of iron reduction that were significantly higher than cells containing an empty vector. *<u>These findings show that BiCASP can be used to validate the EET presented by individual MtrA mutants discovered when screening a combinatorial library of mutants</u>*.

### Implications

The screening of the MtrA mutant library using BiCASP illustrates how this electrochemical HTS platform will transform the engineering of proteins that regulate the electrical communication between cells and materials. The data reported herein, which included >160 unique cellular samples, would have taken more than a month of continuous EET monitoring using a conventional BES,^24–28,61^ provided that a 12-channel potentiostat was used to make parallel measurements. Instead, BiCASP only required a couple hours for data collection and a single potentiostat channel. In total, dozens of decaheme cytochrome mutants were discovered using BiCASP that present real-time current exceeding the negative control, which increases the number of MtrA mutants having their direct EET reported by an order of magnitude.^50^ Mapping all insertion variants that were verified using biological replicates reveals that MtrA retains EET following peptide insertion at backbone sites that are on both the periplasmic and extracellular faces (Figure 5E). Furthermore, the consistency of BiCASP results with the established ferric iron assay supports its accuracy, while offering advantages in the speed of measurement and avoiding the need for anaerobic sample preparation.^62^ When screening for electroactive cells, BiCASP data is preferred over iron reduction because BiCASP directly monitors real-time current at the cell surface under the conditions where cells are used as components in bioelectronics.^16^

By addressing limitations of traditional BESs, BiCASP offers new possibilities for high-throughput screening of electroactive microbes. As illustrated herein, BiCASP represents a powerful tool for genetic screening, which can be used to advance our fundamental understanding of EET mechanisms. By coupling real time measurements of cellular current with genetic tools that knock down the expression of specific proteins,^63^ BiCASP can help elucidate pathways that mediate EET in diverse microbes beyond the MtrA pathway.^31,64^ Additionally, by expressing libraries of mutant protein electron carriers, BiCASP can be used as a screening tool to understand the molecular recognition of the proteins that support electron transfer required for EET,^31^ including periplasmic electron carriers.^65^ BiCASP can also be used to engineer proteins that exhibit conditional electron transfer activity.^36–39^ Such proteins are currently needed to enable the creation of the next generation of bioelectronic devices, such as digital pills,^66,67^ that use microbes to report on environmental chemicals and materials.^17^ Finally, BiCASP can be used for fundamental microbiology studies that seek to understand the prevalence of EET in different microbiomes and across different growth conditions. As an example, wastewater treatment plants are rich in families of organisms that contain the Mtr pathway,^32,33^ including Aeromonadaceae and Pseudomonadaceae,^68^ although the EET for most of these organisms remain uncharacterized.

While this work demonstrates the utility of BiCASP for HTS of MtrA mutants in electroactive microbes, the potential applications of this platform extend far beyond this immediate need. The ability to perform rapid, parallelized electrochemical measurements in individually addressable microwells makes BiCASP a versatile tool for a broad range of biological systems. For example, electrochemical HTS strategies enabled by BiCASP can be applied to monitor metabolic activity,^69^ redox signaling,^70^ and ion flux in mammalian cells,^11^ enabling drug screening, toxicity profiling, and studies of tumor cell heterogeneity. The label-free, miniaturized, and real-time nature of electrochemical detection also makes it compatible with organ-on-chip models and personalized medicine approaches.^71–73^ Thus, BiCASP represents a foundational platform with the potential to accelerate discovery in both prokaryotic and eukaryotic systems, advancing the frontiers of bioelectronics, cellular engineering, synthetic biology, and therapeutic development.

## Methods

### Materials

Acrylic plates for fabricating the bottom wells (1/4^th^ inch; DUYKQEM), top wells (1/16^th^ inch; Acrylic Mega Store), and base plates (1/16^th^ inch) were purchased from McMaster-Carr (Boston), DUYKQEM, and Acrylic mega store. Carbon felt (1/8^th^ inch) was purchased from ThermoFisher Scientific. Carbon ink used for the CE and WE and silver-silver chloride ink (60:40) used for the RE were from Kayaku Advanced Materials. Polyethylene terephthalate sheets, Luria Bertani (LB) agar, LB broth (Miller), M9 minimal medium, kanamycin sulphate, and sodium lactate were purchased from Sigma Aldrich. All other chemicals and materials used for biological screening were from Fisher Scientific, VWR, Millipore Sigma, Merck, Grainger, Research Products International, and Avantor.

### BiCASP fabrication

The electrodes and wells for the BiCASP, which were designed using Adobe Illustrator, have four components, designated A to D. (Component A) The bottom electrodes, which include the CE and RE, consist of parallel lines per row (2 mm for CE and 1 mm for RE) separated by 2.8 mm with connection pads located at opposite ends for each electrode. (Component B) The top electrode, or the working electrode (WE), is composed of 2 mm lines and 0.6 cm circles along the line. All three electrodes had a 1.2 cm^2^ connection pad on one end. Stencils for the three electrodes were cut from polyethylene terephthalate (PET) sheets (Chartpak) using either a Universal Laser System (ULS) with a 10.6 µm CO₂ laser or a Speedy 360 Trotec Ruby laser cutter. The CE/RE and WE stencils measured 18.5 × 9 cm and 11 × 13.5 cm, respectively. After cutting, the stencils were affixed to another new PET sheet measuring 21.6 × 28 cm. The CE and WE were screen-printed using conductive carbon ink, while the RE was screen-printed with silver-silver chloride ink. The screen-printed PET sheets were then placed in an oven at 100°C for 5 minutes to facilitate the evaporation of the solvent in the ink and ensure complete drying. Lower temperatures could be used (85°C), but these required longer treatment periods (1 hour). This procedure was repeated twice. Once the inks had completely dried, the stencils were peeled off, leaving behind defined electrodes on the underlying PET sheet(s). Sterilized carbon felt was cut using a 0.6 cm hole cutter, following pretreatment with a UVO cleaner (Jelight Company Inc.) for 20 minutes, and affixed to the WE using carbon ink. To ensure the solvent in the ink evaporated, the material was heat-treated. (Component C) Acrylic wells, consisting of square wells measuring 0.6 by 0.6 cm, were cut from 1/4^th^ and 1/16^th^ inch acrylic sheets (13.5 x 9 cm), serving as wells for the bottom and top electrodes, respectively. (Component D) To support the top and bottom PET electrode, acrylic base plates, comprised of rectangular plates (8 x 6 cm) cut from a 1/16^th^ inch acrylic sheet, were generated. Each component featured four holes at the corners to facilitate proper alignment of all device parts.

### BiCASP assembly and operation

The CE/RE and WE on the PET sheets were affixed to separate acrylic base plates using a waterproof silicone adhesive sealant (Permatex). A thick acrylic well (1/4^th^ inch), was adhered to the bottom electrode with a substantial layer of silicone adhesive to prevent leakage. Similarly, a thinner acrylic well (1/16^th^ inch) was attached to the top electrode. The top electrode, comprising the base plate, PET electrode, and carbon felt, underwent plasma treatment in a Plasma chamber for 10 to 20 minutes to render the carbon felt hydrophilic. The capacity of each microwell is about 500 µL. For all experiments, 400 µL of the liquid medium, either M9 or SBM plus the lactate carbon source, was added to the bottom acrylic well, while 50 µL of the bacterial solution was applied directly to the carbon felt. The top electrode was then placed on the bottom electrode to close and loosely seal the device. Electrode wires from the well of interest were connected to a potentiostat (CHI workstation) through alligator wire and clips through the external connection pad on the BiCASP. As a quality control for fabrication, the diffusion of food dye between wells was monitored for one hour, and each electrode was tested for continuity using a multimeter. When devices were reused, the carbon felt pads were removed, the device was washed with water, new carbon felt pads were attached with carbon ink, and the device was dried for 20 minutes at 85°C. The electrochemical properties were determined by cyclic voltammetry (CV) and chronoamperometry (CA).

### Strains and plasmids

*S. oneidensis* MR-1 was from the American Type Culture Collection (ATCC 700550), while the strain lacking the MtrA gene (*S. oneidensis* JG665), which has the genotype *ΔmtrA/ΔmtrD/ΔdmsE/ΔSO4360/ΔcctA*,^53^ was kindly provided by Jeff Gralnick. The plasmid used for complementation of JG665 (pJA018) expresses MtrA using a TetR promoter,^74^ while the empty vector (pIC004) used as a negative control lacks the MtrA gene.^50^ These plasmids both contain an RSF1010 origin of replication and confer kanamycin resistance. The library of MtrA mutants was previously introduced into pJA018,^50^ such that they contain randomly inserted sequences that code for the octapeptide SGRPGSLS. All plasmids screened or selected from this library were sequenced using Oxford Nanopore (Plasmidsaurus, Inc.). The sequences of the MtrA variants that were characterized are provided as supporting information.

### Growth medium

Cells were grown in either LB or Shewanella Basal Medium (SBM).^53^ SBM contains minerals (7.5 mg/L N(CH₂CO₂H)₃, 0.5 mg/L MnCl₂·4H₂O, 1.5 mg/L FeSO₄·7H₂O, 0.85 mg/L CoCl₂·6H₂O, 0.5 mg/L ZnCl₂, 0.2 mg/L CuSO₄.5H₂O, 0.025 mg/L KAl(SO_4_)_2_·12H_2_O, 0.025 mg/L H₃BO₃, 0.45 mg/L Na₂MoO₄, 0.5 mg/L NiCl₂, 0.1 mg/L Na₂WO₄·2H₂O, and 0.5 mg/L Na₂SeO₄), vitamins (0.01 mg/L biotin, 0.01 mg/L folic acid, 0.1 mg/L pyridoxine HCl, 0.025 mg/L thiamine, 0.025 mg/L nicotinic acid, 0.025 mg/L pantothenic acid, 0.5 µg/L vitamin B12, 0.025 mg/L p-aminobenzoic acid, and 0.025 mg/L lipoic acid), salts (8.6 mM NH₄Cl, 1.3 mM K_2_HPO_4_, 1.65 mM KH₂PO₄, 475 µM MgSO4·7H_2_O, and 1.7 mM (NH₄)₂SO₄), HEPES (100 mM), lactate (20 mM), and casamino acids (0.05% w/v). In cases where kanamycin (50 µg/mL) was added to select for plasmids, the pH was adjusted to 7.2 using NaOH, and the medium was sterilized using a 0.22 µm filter.

### Cyclic voltammetry and signal optimization

Cyclic voltammetry was performed using 1 mM ferricyanide/ferrocyanide redox medium prepared in 0.10 M KCl. The potential sweeping window was set from −0.2 to 0.4 V (vs. Ag/AgCl reference electrode) with a scan rate of 1 mV/s. Cellular measurements used *Shewanella oneidensis* MR-1. To generate cultures for analysis, single colonies from LB agar plates were used to grow 5 mL LB cultures. Following incubation at 30°C overnight while shaking at 250 rpm, an aliquot (500 µL) was used to inoculate a 50 mL LB culture. After growing at 30°C for 18 hours while shaking (250 rpm), cells were then harvested by centrifugation, washed using M9 minimal medium, and resuspended in M9 medium to the indicated optical densities.

### Screening the MtrA Mutant Library

The MtrA library was screened using two approaches.^50^ First, to enrich for cells expressing MtrA mutants that support EET, cells transformed with the vector library were diluted to an optical density of 0.1 in 100 mL of SBM and grown in crimp top serum bottles (Millipore Sigma) containing 30 mM ferric citrate under anaerobic conditions for 24 hours. Following 24 hours at 30°C, while shaking at 250 rpm, cells were diluted 10-fold, and an aliquot (50 µL) was spread on LB-agar medium containing kanamycin (50 µg/mL). Individual colonies observed after 24 hours were used to inoculate SBM cultures (5 mL) in 24 deep-well plates, which were grown at 30°C for 48 hours while shaking at 250 rpm. The resulting cultures were analyzed for EET using BiCASP and an iron reduction assay.^62^ Second, to determine if BiCASP can identify EET-active cells without enriching prior to analysis, cells transformed with the vector library were plated directly onto LB-agar medium containing kanamycin without a preculturing step. After incubating at 30°C for 24 hours, individual colonies (n = 40) were used to inoculate SBM medium (2.5 mL). After growing aerobically at 30°C for 48 hours while shaking at 250 rpm, cultures were used to inoculate fresh SBM cultures containing antibiotic (5 mL). After 48 hours, each culture was evaluated using BiCASP and the iron reduction assay. To evaluate EET, liquid cultures were pelleted by centrifugation at 5,000 rpm (4,472 xg) for 10 minutes and resuspended with SBM at an OD_600_ of 20. This protocol typically yielded ∼650 µL of concentrated cells, 50 µL of which was applied directly to a BiCASP working electrode. In parallel, each well was filled with SBM (400 µL) containing sodium lactate (100 mM). Devices were immediately assembled, and current was measured within 2 minutes of assembling using working electrodes poised at +0.2 V using a VMP-300 BioLogic potentiostat. When quantifying microbial current, signals were evaluated by comparing the current at the sixty-second time point.

### Plasmid sequencing

To establish the MtrA variants expressed in each culture, plasmids were purified (Qiagen Miniprep) and subjected to Oxford nanopore sequencing. Sequencing revealed that 7.5% of the samples had either a large sequence deletion in the MtrA gene (n = 2) or a mixture of plasmids (n = 1). The currents and iron reduction rates from these samples are not shown. Variants were numbered as previously described using the native residue that immediately follows the peptide insertion.^50^

### Iron Reduction Assay

Whole cell iron reduction was performed as described.^62^ Within shallow 96-well plates (Corning 96-Well, Cell Culture-Treated, Flat-Bottom Microplate, Fisher Scientific), cultures used for BiCASP measurements (OD = 20 at 600 nm) were diluted 50-fold into fresh SBM (100 μL) supplemented with 5 mM ferric citrate and ferrozine (1 mg/mL) under aerobic conditions. Biological replicates performed using the mutants screened from libraries were cultured overnight in LB (1 mL) and diluted similarly in SBM (1 mL) prior to performing the assay. After diluting cells 50-fold into SBM, plates were transferred to an anaerobic chamber (Vacuum Atmospheres), a transparent seal was applied over the plate, edges were lined with silicone grease, and lids were used to seal plates. Absorbance (562 nm) was measured every 3 minutes using a TECAN Spark plate reader at 30°C with double orbital shaking at 90 rpm and 3 mm amplitude. After subtracting off the initial absorbance values at time zero, the resulting data was fit to an exponential model:

> Fe(II) = K·(exp(μt) − 1)

where *Fe(II)* represents the absorbance signal, *t* is time in hours, *K* is a constant representing the initial number of cells used to inoculate the assay culture, and *μ* represents the rate constant.

### Structural visualization

The crystal structure of the Mtr complex from *Shewanella baltica* (PDB 6R2Q) was utilized for structural analysis.^75^ Insertion sites were mapped onto the structure using the PyMOL Molecular Graphics System, Version 2.5.2 (Schrödinger, LLC). Additional molecular graphics were generated using UCSF ChimeraX, developed by the Resource for Biocomputing, Visualization, and Informatics at the University of California, San Francisco, with support from National Institutes of Health R01-GM129325 and the Office of Cyber Infrastructure and Computational Biology, National Institute of Allergy and Infectious Diseases.

### Statistical analysis

For BiCASP data, signal optimization data were analyzed using linear regression analysis (a = 0.05), while strain and variant comparisons were performed using one or two-tailed unpaired Welch’s t test (a = 0.05). For the Fe(III) reduction assay, data was first fit to the exponential model to obtain rates, and then rates were compared using a two-tailed unpaired Welch’s t test.

## Supplemental material

Supplemental Figure S1-S5 are provided as a single PDF file. Also, the plasmids identified in library screening are provided as a single XLSX file.

## Resource availability

### Lead contact

Requests for further information and resources should be directed to and will be fulfilled by the lead contact, Sameer Sonkusale

### Materials availability

The plasmids generated in this study have been deposited in Addgene.

### Data and code availability

Adobe Illustrator files (.ai); 2D design for all parts of a 4x6 BiCASP (Component A-D)

Python script files (.py); microcontroller firmware, and Python codes to drive the GUI

Multiplexer design files (.sch); schematic and circuitry

Gerber file (.GBR); design files and layout information

Application file (.exe); packaged folder to execute the MUX

## Supporting information

Supplemental section

Plasmids identified in library screeening

## Acknowledgments

Research was also sponsored by the Army Research Office and was accomplished under Grant Number W911NF-22-1-0239. The views and conclusions contained in this document are those of the authors and should not be interpreted as representing the official policies, either expressed or implied, of the Army Research Office or the U.S. Government. The U.S. Government is authorized to reproduce and distribute reprints for Government purposes, notwithstanding any copyright notation herein. We are also grateful for support from the National Science Foundation grant 2223678 and the Office of Basic Energy Sciences of the U.S. Department of Energy grant DE-SC0014462. The authors would like to thank Prof. Caroline M. Ajo-Franklin for graciously donating *Shewanella oneidensis* (MR-1) and for her guidance in setting the culturing protocol at Tufts university.

## Author contributions

H.S., J.J.S., and S.S.: Conceptualization;

H.S., P.B., K.S., S.V.D., A.T., C.W., and R.P.: Investigation;

H.S., P.B., K.S., S.V.D., C.A., A.T., C.W., R.P., A.S., M.D.C., S.B.: Methodology;

H.S., P.B., K.S., and A.T.: Data curation;

H.S., P.B., and A.T.: Formal analysis;

H.S., J.J.S., and P.B.: Writing – original draft;

All authors: Writing, review & editing;

H.S. and P.B.: Visualization;

S.V.D., C.A., and C.W.: Software;

J.J.S. and S.S.: Resources;

J.J.S., and S.S.: Supervision;

J.J.S., R.V., and S.S.: Project administration;

J.J.S., R.V., and S.S.: Funding acquisition.

