## Supplemental section for "A Bioelectrochemical Crossbar Architecture Screening Platform (BiCASP) for Extracellular Electron Transfer"

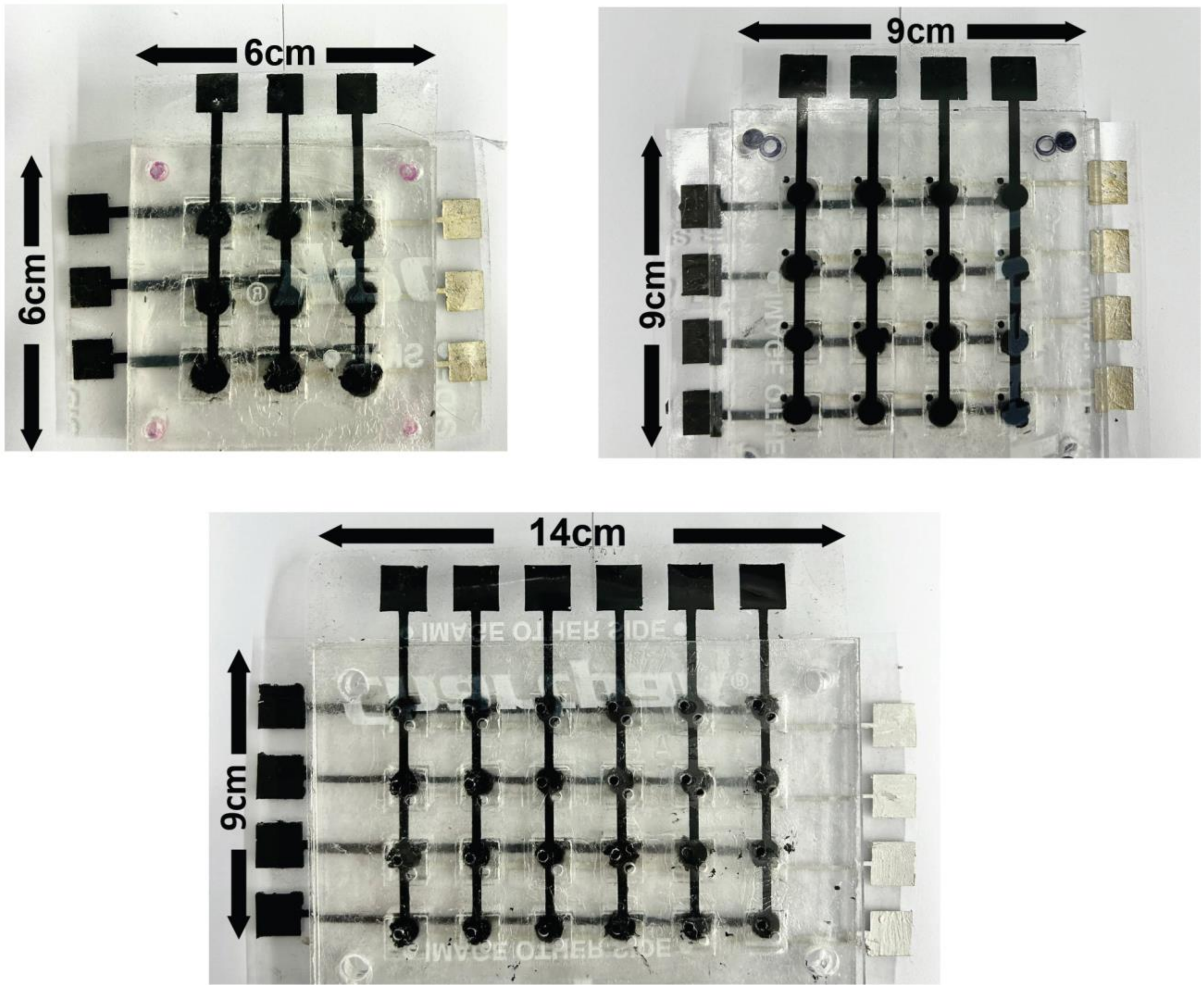


**Figure S1. BiCASP fabricated with varying numbers of wells.** BiCASP devices can be fabricated with varying numbers of wells that leverage the crossbar architecture. While 9-, 16-, and 24-well formats are shown, this approach can be used to create more complex formats such as 96- and 384-well formats to enable HTS biological applications.

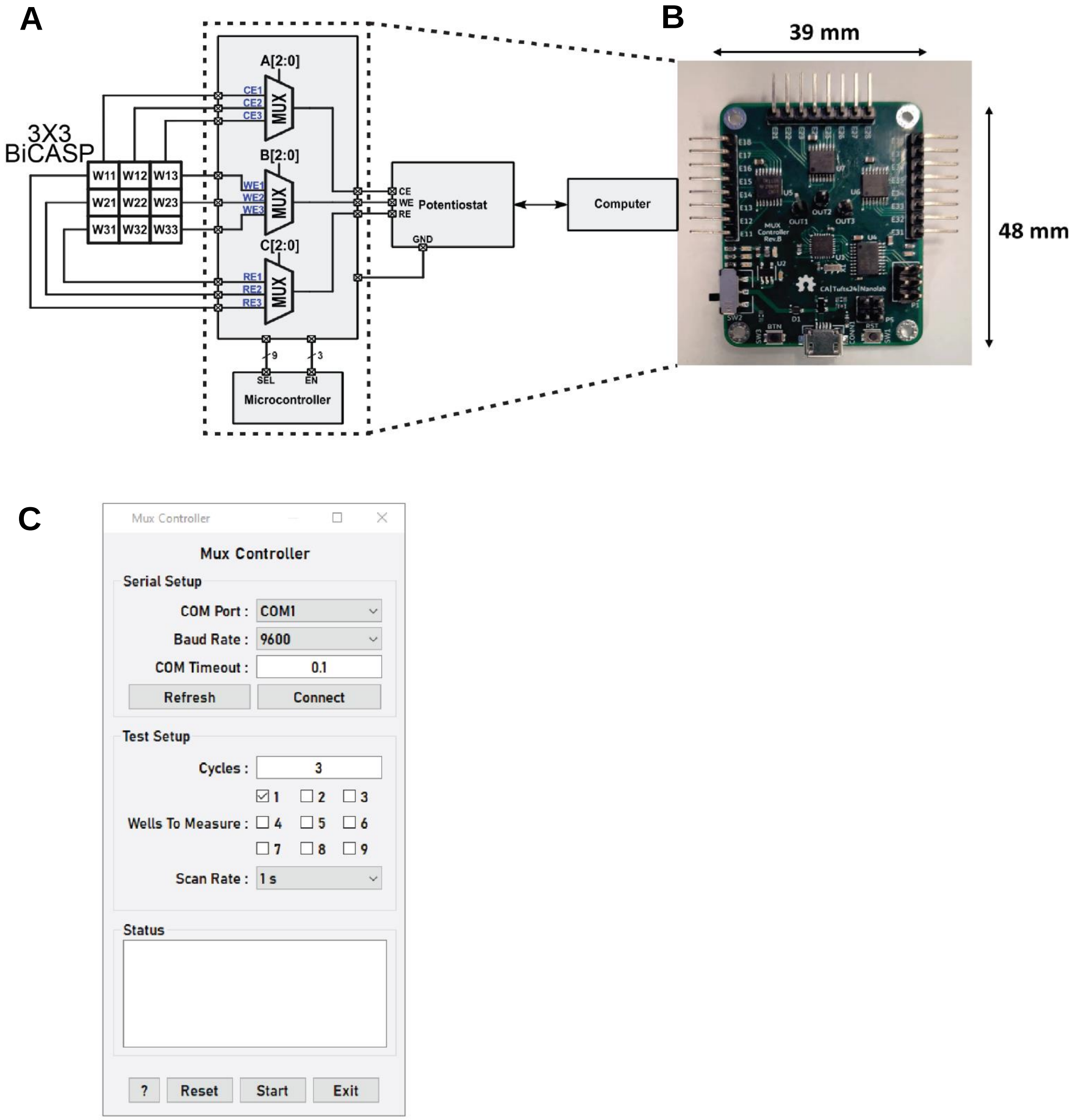


**Figure S2. Integration of an electrical multiplexer (MUX) with the BiCASP. (A)** Block diagram of the MUX showing all the connections between the BiCASP, microcontroller and the potentiostat for a 3x3 well array. **(B)** Image of the printed circuit board (PCB) with precision multiplexer (TMUX 1108), microcontroller (ATmega328p), UART bridge for PCB to computer connection, 8x3 input lines with three output lines. **(C)** Graphics user interface (GUI) to input the multiplexing parameters for switching between the wells

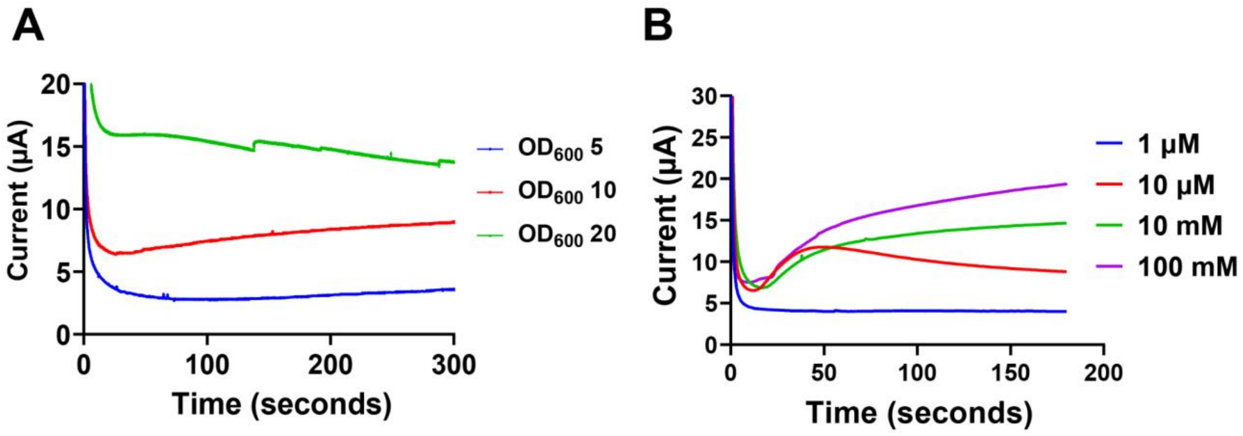


**Figure S3. Identifying BiCASP conditions that yield strong biological signals.** Chronoamperometric signal showing direct electron transfer measurement for different concentrations of (**A**) *S. oneidensis* MR-1 (including $OD_{600}$ 5, 10 and 20) and (**B**) lactate (ranging from 1 μM to 100 mM). All measurements were performed in different wells at different times. Data represent average of three biological replicates.

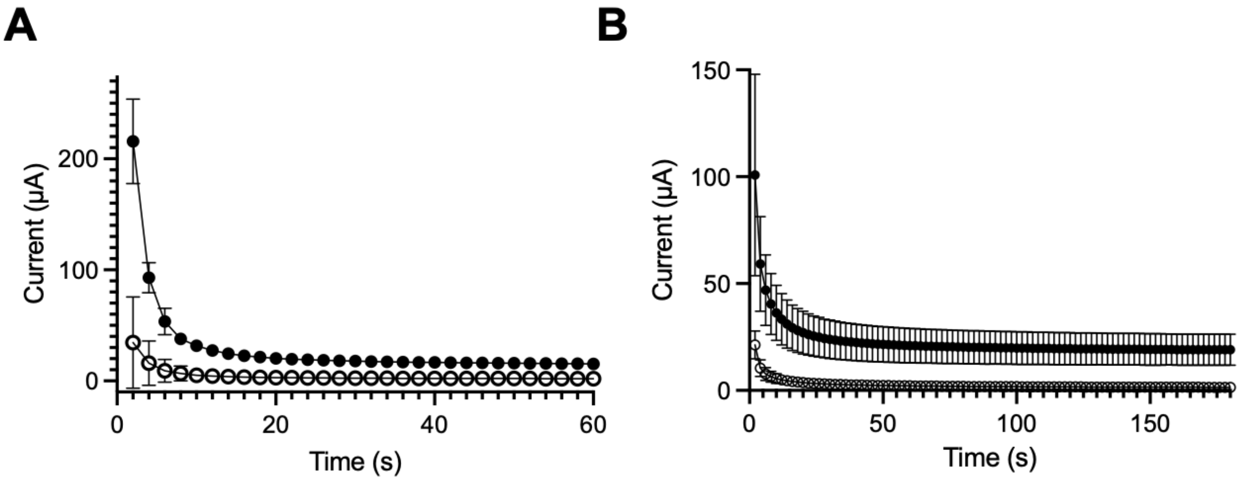


**Figure S4. BiCASP current stabilizes in less than a minute.** The first three minutes of data collected for (**A**) *S. oneidensis* MR-1 (closed circles) and JG665 (open circles) reveals that the signals converge on a stable signal in less than a minute. Data represent the average of twelve biological replicates with error bars representing one standard deviation. (**B**) *S. oneidensis* MR-1 transformed with a vector that expresses MtrA (closed circles) also presents a dynamic signal that stabilizes in less than a minute and is greater than cells transformed with an empty vector (open circles). With these measurements, the data represents the average of eight biological replicates with error bars representing ±1σ.

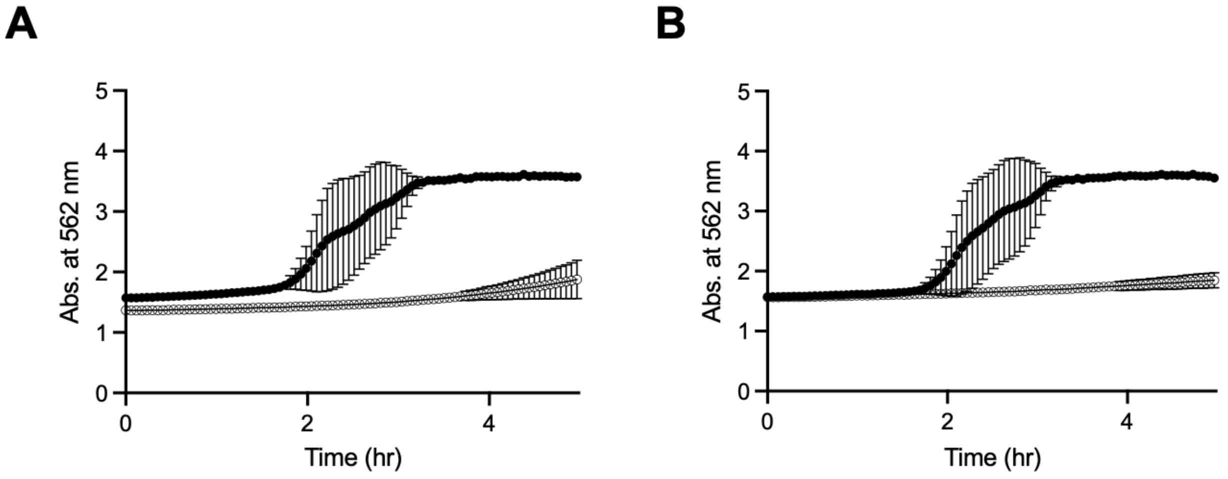


**Figure S5. The iron reduction assay requires longer duration data collection.** With the iron reduction assay, reduced iron is visualized by continuously monitoring absorbance at 562 nm. This assay was used to compare EET mediated (**A**) *S. oneidensis* MR-1 (solid circles) and *S. oneidensis* JG665 (open circles) and (**B**) *S. oneidensis* JG665 transformed with an empty vector (open circles) and a vector that constitutively expresses MtrA (solid circles). The data shown represents the average of four biological replicates with error bars representing one standard deviation.
